# Signaling through the Dystrophin Glycoprotein Complex affects the stress-dependent transcriptome in Drosophila

**DOI:** 10.1101/2022.06.15.496303

**Authors:** Travis D Carney, Rucha Y Hebalkar, Evgeniia Edeleva, Ibrahim Ömer Çiçek, Halyna R Shcherbata

## Abstract

The Dystrophin Glycoprotein Complex (DGC) is a cell membrane-spanning complex that links the extracellular matrix with the intracellular cytoskeleton. Deficiencies in the DGC in humans cause muscular dystrophies (MDs), a group of inherited, incurable disorders associated with heterogeneous muscle, brain, and eye anomalies. To advance disease diagnostics and develop new treatment strategies, it is essential to understand the genetic pathways that are perturbed by DGC mutations and the mechanisms underlying these pathologies. Stresses such as nutrient deprivation and aging cause a reduction of muscle mass can be exacerbated by a reduced content of the DGC in membranes, whose integrity is vital for muscle health and function. This illustrates that the DGC plays a role in stress-response pathways. Therefore, it is important to investigate the influence of stress not only on healthy individuals but also on the wellbeing of MD patients. Moreover, the DGC has also emerged as an integral component in multiple signaling pathways, demonstrating an important yet poorly understood connection between intercellular forces and regulation of gene expression and illustrating the importance of understanding DGC-related transcriptional effects. Here, we utilize a Drosophila model to investigate the transcriptomic changes in mutants of four different DGC components under unstressed, temperature-stressed, and starvation-stressed conditions. Our analysis reveals a group of genes that exhibit DGC-dependent gene regulation. We identify large groups of genes that are differentially regulated in response to either temperature or starvation stress. Importantly, we also identify groups of genes with expression patterns dependent on the DGC signaling pathway for a proper stress response. This work reveals a novel function of the DGC in stress-response signaling. The view of the DGC as a regulatory unit involved in the stress response will give new insights into the etiology of symptoms of MDs and possible directions of symptomatic treatment and relief, and it will ultimately aid in a better understanding of DGC signaling and regulation under normal and stress conditions.

## Introduction

The Dystrophin Glycoprotein Complex (DGC) is a cell membrane-associated protein complex that connects the extracellular matrix (ECM) to the cytoskeleton of the cell. The core components of the DGC are the transmembrane, ECM-associated protein Dystroglycan (Dg); cytoplasmic Dystrophin (Dys); and cytoplasmic Syntrophin (Syn) proteins (Fig. 1A). DGC dysfunction is associated with a group of diseases, the muscular dystrophies (MDs), which have deleterious and sometimes fatal effects on muscles and the nervous system. For example, loss of Dys results in Duchenne MD (DMD) (Ervasti & Campbell, 1993), and aberrant glycosylation of Dg leads to severe forms of congenital and late onset MDs (Barresi *et al*, 2004; Bigotti & Brancaccio, 2021; Brun *et al*, 2018; Goddeeris *et al*, 2013; Imae *et al*, 2021; Joseph & Campbell, 2021; Mantuano *et al*, 2021; Munot *et al*, 2022; Xie *et al*, 2022; Yatsenko *et al*, 2021). In contractile muscle cells, DGC-dependent linkage of the ECM and intracellular cytoskeleton accounts for the stability and mechanical stress resistance of the sarcolemma, limiting contraction-initiated damage (Demontis *et al*, 2013; Gao & McNally, 2015; Larsson *et al*, 2019). MD patients experience progressive muscle degeneration and often die because of heart or respiratory failure. DMD patients also suffer from cognitive impairment (Perronnet & Vaillend, 2010), and some patients with congenital MDs exhibit structural brain abnormalities, mental retardation, abnormal neuronal migration, and white matter changes (Cohn, 2005). In addition to muscular and neurological deficits, in mice it has been reported that DGC deficits can result in reduced male fertility (Chen *et al*, 2017; Hernandez-Gonzalez *et al*, 2005), and in Drosophila, these deficits cause systemic problems such as an inability to maintain temperature homeostasis (Takeuchi *et al*, 2009).

**Figure 1:**
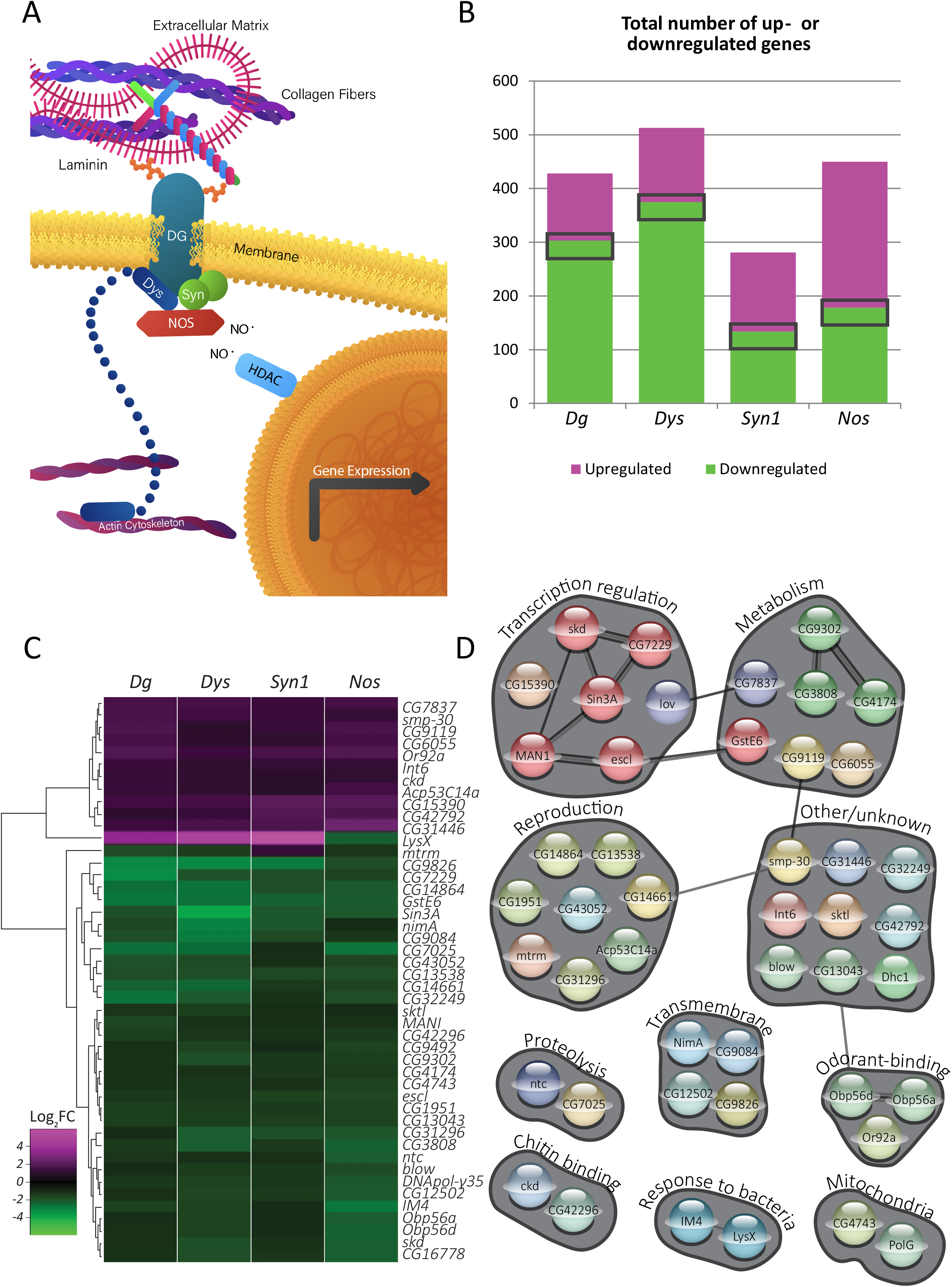
DGC-dependent signaling exerts a defined transcriptional response. A. Schematic of the DGC and associated proteins. The transmembrane protein Dystroglycan (Dg) is the core component of the complex. Its glycosylated extracellular domain associates with the extracellular matrix, while its intracellular portion associates with Dystrophin and Syntrophin (Dys and Syn), which can act as structural and signaling scaffolds, attaching to the intracellular cytoskeleton as well as to signaling molecules such as nitric oxide synthase (Nos). The nitric oxide (NO) produced by Dys-associated Nos can nitrosylate intracellular proteins such as histone deacetylases (HDACs), affecting the expression of downstream genes. B. Numbers of genes found to be dysregulated in flies mutant for each of four DGC components: *Dg* (*DgO55/DgO86*); *Dys* (*Df(3R)Exel6184*); *Syn1* (*ΔSyn15-2/Df(3L)BSC450C*); *and Nos* (*ΔNos*^*15*^*/Df(2L)BSC230*). Numbers of dysregulated genes ranged from 281 for *Syn1* to 513 for *Dys*. The gray rectangles represent the 46 genes that are dysregulated in all four mutants. C. Heat map illustrating the 46 genes that are dysregulated in all four mutants assayed. 11 genes are upregulated and 33 downregulated in all mutants; only 2 genes, *LysX* and *mtrm*, are differentially regulated depending on genotype. D. STRING-based clustering of gene products of the 46 genes that are dysregulated in all four DGC mutants. Clusters are labeled according to the functions and processes in which the genes are involved. Colored lines indicate putative and experimentally determined physical interactions.

The DGC is also involved in cell signaling. Syntrophins have multiple protein-protein interaction motifs and can serve as adaptor proteins capable of binding to heterotrimeric G proteins and neuronal nitric oxide synthase (nNos), among other signaling modules (Cacchiarelli *et al*, 2010; Xiong *et al*, 2009; Zhou *et al*, 2006). Syntrophins comprise a cytoplasmic platform to which nNos can bind and produce nitric oxide (NO), an important signaling molecule that acts via the nitrosylation of intracellular proteins. This nitrosylation serves to inhibit mammalian histone deacetylase 2 (HDAC2), leading to activation of HDAC2-responsive genes, including microRNAs necessary for the differentiation of muscle progenitor cells (Cacchiarelli *et al*., 2010). In flies, Dg and Nos signal cooperatively via a feedback loop in which Nos activity promotes the expression of a cluster of microRNAs that directly target the *Dg* transcript. Consistent with a functional DGC providing a signaling platform for Nos, overexpression of Dg results in increased production of NO by Nos (Yatsenko *et al*, 2014b). As the DGC is a mechanosensory complex, this conserved link demonstrates an important connection between intercellular forces and transcriptional activity.

Another important example of the link between the DGC, signaling, and downstream transcriptional activity is demonstrated by the relationship between the DGC and Hippo signaling in both mice and Drosophila. In murine cardiomyocytes, the phosphorylated effector of Hippo signaling, Yap, is directly bound by the Dg ortholog Dag1, and thus sequestered by the membrane-associated complex. This sequestration serves as an important control on cardiomyocyte proliferation, evidenced by an overproliferation phenotype at the site of cardiac injury in Hippo-DAG1 double mutants (Morikawa *et al*, 2017; Vita *et al*, 2018). Similarly in flies, the Hippo effector Yorkie physically associates with the DGC, as does another component of the signaling cascade, Kibra. These interactions were shown to promote the maintenance of muscle integrity during aging in adult flies (Yatsenko *et al*, 2020).

A fundamental factor affecting organismal physiology is temperature. Genomics studies in various model organisms, such as mice, flies, worms, and yeast have demonstrated that in response to heat stress, a rapid and transient reprioritization of the gene expression program occurs. These changes include repression of genes involved in growth and cell proliferation, rearrangement of DNA and chromatin, regulation of energy metabolism and redox state of the cells, alternative splicing, and proteostasis (Evans, 2015; Gasch *et al*, 2000; Lopez-Maury *et al*, 2008; Sorensen *et al*, 2016). Exposure to high ambient temperatures can result in high morbidity and mortality (Shindell *et al*, 2020; Vicedo-Cabrera *et al*, 2021). Extreme or prolonged heat can overwhelm thermoregulatory capacity even in healthy persons, but it is especially dangerous for patients with muscle disorders (Cheshire, 2016). For example, patients with muscular dystrophies have a high risk of malignant hyperthermia and heart failure as a response to anesthetic agents (Hayes *et al*, 2008; Ohkoshi *et al*, 1995; Rohde *et al*, 2014). In these individuals, a drastic increase of Ca^2+^ in skeletal muscle leads to sustained contractions, heat generation, and a dangerous increase in body temperature. In Drosophila larvae, Dg mutation also causes increased intracellular Ca^2+^ concentration and concomitant oxidative metabolism, resulting in a cold-seeking behavior that is rescued by the transgenic re-introduction of Dg (Takeuchi, et al. 2009). However, the role of the DGC in thermoregulation remains elusive. Moreover, since increasing worldwide environmental temperatures have dire health effects, particularly among urban dwellers and people with metabolic and cardiopulmonary disorders, (Burkart *et al*, 2021; Ebi *et al*, 2021; Shindell *et al*., 2020; Vicedo-Cabrera *et al*., 2021), it is important to understand the influence of heightened temperature not only on patients with muscular dystrophies but also on the wellbeing of healthy individuals.

In addition to genetic disorders (muscular dystrophies), other physiological and pathological stimuli (*e*.*g*., fasting and cachexia) can cause muscle wasting. Starvation usually results in muscle atrophy, which is loss of muscle mass due to an increase in protein degradation or a decrease in protein synthesis (Piccirillo *et al*, 2014). It is an integral feature of systemic diseases including cancer, cachexia, cardiac failure, AIDS, and sepsis. One important aspect of the stress response to dietary restrictions is an alteration in muscle metabolism that leads to the decreased usage of carbohydrates, so they can be spared for the organs and tissues where glucose is essential, such as the central nervous system. Loss of muscle mass due to aging, also known as sarcopenia, is often associated with muscle disuse, fasting, extrinsic changes in innervation, stem cell function, and endocrine regulation of muscle homeostasis (Demontis *et al*., 2013). This loss of muscle mass is triggered, in part, by the reduced content of the DGC in membranes, whose integrity is vital for muscle health and function. In fact, the reduction in DGC content seems to precede and promote age-associated muscle atrophy and associated weakness and frailty (Bengtsson *et al*, 2022). Conversely, stabilization of the DGC on the muscle membrane markedly attenuates atrophy (Eid Mutlak *et al*, 2020). Therefore, understanding the role of the DGC in muscle maintenance upon metabolic stress is extremely important for understanding the molecular mechanisms that cause muscle atrophy during aging and in various catabolic states (*e*.*g*. starvation, type-2 diabetes). However, it is not clear how this mechanosignaling complex affects various components of the cellular stress response.

The components of the DGC are evolutionarily conserved from Drosophila to mammals but exist in Drosophila with significantly less redundancy (Greener & Roberts, 2000). As in mammals, the DGC components in Drosophila are expressed not only in muscle but also in nervous and other tissues (Bogdanik *et al*, 2008; Dekkers *et al*, 2004; Marrone *et al*, 2011a; van der Plas *et al*, 2006). Flies deficient for Dys or Dg develop phenotypes similar to MD patients, both in the muscle and in the nervous systems. They experience a shortened lifespan, decreased mobility, age-dependent muscle degeneration, and defective neuron differentiation (Shcherbata *et al*, 2007). Using *Drosophila melanogaster* as a MD model, our lab previously demonstrated that temperature, metabolic stress, oxidative stress, and aging can enhance muscle degeneration in flies mutant for *Dys* or *Dg* and can promote degeneration even in wild type flies (Kucherenko *et al*, 2011). In addition, we found a group of both dystrophy- and stress-dependent microRNAs that are up- or downregulated due to high-temperature stress in wild type flies but fail to change their expression levels under high temperature in dystrophic flies (Marrone *et al*, 2012). These data support a concept that there is a signaling pathway under stress between the cell membrane-associated DGC and nuclear gene expression.

To find novel DGC-dependent genetic pathways and to further investigate the relationship between the DGC and transcriptional stress response, we performed RNA-sequencing analysis from whole flies under unstressed conditions, under temperature stress, and under starvation stress. We compared control flies to mutants of four different DGC components: Dg, Dys, Syn1, and Nos (Supp. Table 1). From this analysis, we uncovered a group of genes that are dysregulated in all four DGC mutants in unstressed conditions, representing genes that are specifically regulated by DGC-dependent signaling mechanisms. Consistent with the DGC comprising a coherent signaling center, nearly all of these genes were similarly regulated (up- or down-regulated) in all four mutants. By comparing stressed to unstressed control flies, we identified several hundred genes that are differentially regulated in response to either temperature or starvation stress. Consistent with previous studies (Lecheta *et al*, 2020; Sorensen *et al*, 2005; Telonis-Scott *et al*, 2013; Zhou *et al*, 2012), we found that the majority of the dysregulated genes are downregulated by temperature stress, but more genes are upregulated by starvation. This pattern is also evident in aged flies and those subjected to oxidative stress (Landis *et al*, 2004). Strikingly, only a small minority of genes are commonly dysregulated by both stresses, illustrating the distinct transcriptional responses to temperature and starvation. Finally, we identified sets of genes whose differential expression patterns under stress are perturbed in mutants with a non-functional DGC, constituting groups of genes dependent on the DGC signaling pathway for a proper stress response. A whole-genome overview of organismal transcriptional changes is an important descriptive analysis of the DGC-dependent stress response. This work reveals a novel function of the DGC in stress-response signaling. The view of the DGC as a regulatory unit involved in the stress response will give new insights into the etiology of MDs symptoms and possible directions of symptomatic treatment and relief.

## Results and Discussion

### DGC-dependent signaling exerts a defined and consistent transcriptional response

To validate that the DGC complex exerts a coherent transcriptional effect, we first investigated the effects of multiple DGC mutants – *Dg, Dys, Syn1*, and *Nos* – in comparison to control flies (Table 1). By using these four different DGC mutant alleles and by hierarchical clustering of dysregulated genes, we were able to distinguish between genes downstream of the DGC complex as a whole and those that could be affected by individual mutant lesions due to DGC-unrelated functions or pleiotropic effects. By this rationale, if the DGC complex is responsible for distinct and regulated transcriptional outputs, we would expect to see that downstream genes are dysregulated similarly in each of the mutants. If, on the other hand, downstream genes were to exhibit differential dysregulation in each of the mutants, this would indicate that these transcriptional perturbations are not a result of a combined effect of the loss of DGC complex function, but rather unrelated or pleiotropic effects.

**Table 1.**
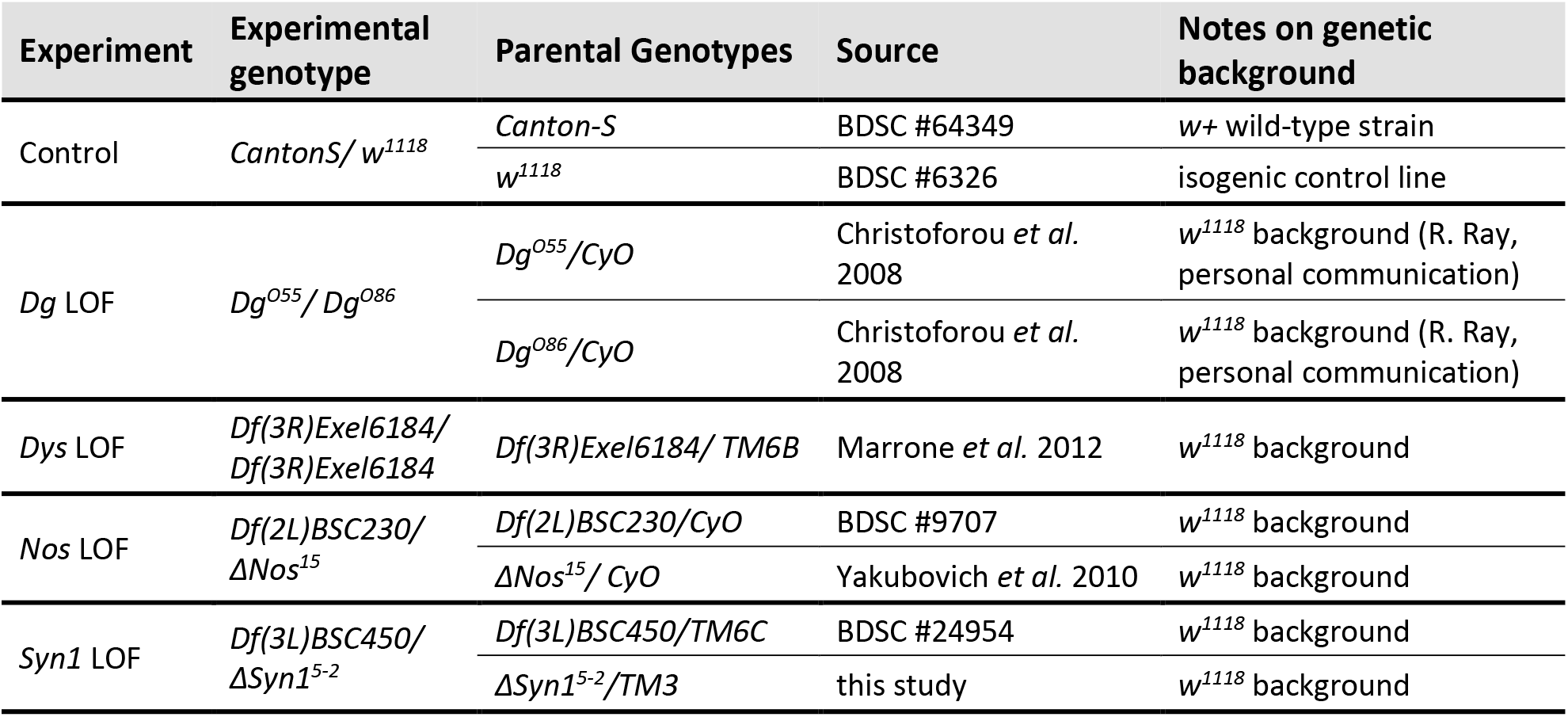

First, we filtered for genes that are up- or downregulated at least 2-fold in any of the DGC mutants when compared with wildtype, irrespective of stress (Fig. 1B). Each of the four DGC mutants exhibits a different number of total dysregulated genes, from 281 in *Syn1* mutants to 513 in *Dys* mutants. The majority of these genes are distinct to each genotype, implying that these are genetic lesion-specific effects. For this reason, we chose to focus instead on the subset of genes that are commonly dysregulated in all four mutant genotypes. With these criteria, we found that 46 genes are at least 2-fold dysregulated in all mutant genotypes (dark rectangles in Fig. 1B; Fig. 1C; Supp. Table 2). Strikingly, 96% (44 of 46 genes) of these genes exhibit a matching up- or downregulation pattern in all DGC mutants. This strongly suggests that the DGC is integral to a defined transcriptional program. Therefore, we conclude that these genes likely represent “DGC-dependent” genes.

As indicated above, each individual DGC lesion results in hundreds of dysregulated genes, while far fewer (46 genes) are commonly dysregulated in all four mutants – the most stringent criterion applied here. Applying the less stringent requirement that genes are dysregulated in only three out of the four mutants, an intermediate number of genes is the result. For example, if we disregard the *Nos* LOF and require only those genes are dysregulated in mutants of the core DGC components *Dg, Dys*, and *Syn1*, we find that an additional 25 genes are dysregulated, or over 50% more genes (Supp. Table 3, genes in bold font). We believe that the core DGC components Dg, Dys, and Syn1 may behave more similarly to one another transcriptionally due to cellular perturbations that result from a physically defective DGC. *Nos* mutation, in contrast, is well known to result in transcriptional changes due to nitrosylation of HDACs, as previously mentioned. Therefore, it is important that we require that genes be dysregulated in the *Nos* mutant as well as in *Dg, Dys*, and *Syn1*, thereby actively selecting for genes that exhibit a transcriptional effect directly downstream of the DGC and increasing our confidence that they can truly be considered DGC-dependent genes (Fig. 1B-1D).

Only two genes, *LysX* and *mtrm*, are differentially regulated across the four mutants. *mtrm* is downregulated in all mutants except *Syn1*, where it is slightly upregulated; the expression changes are much more modest than those exhibited by *LysX. mtrm* expression is highly female-biased and encodes a protein required for proper chromosome segregation during meiosis (Harris *et al*, 2003). As we used male flies for the sequencing experiments, *mtrm* expression level was low in most samples (e.g., *mtrm* was among the lowest expressing 25% of transcripts detected in Canton-S controls and the bottom 5% of *Dg* mutant transcripts).

*LysX* is the strongest upregulated gene in *Dg, Dys*, and *Syn1*, but it is downregulated in the *Nos* mutant (Fig. 1C). This may indicate that *LysX* is regulated by the DGC complex and Nos via independent mechanisms. *LysX* is one of seven lysozyme genes in *Drosophila melanogaster*, all of which are clustered on chromosome 2 near 61F. Lysozymes hydrolyze peptidoglycan in bacterial cell walls, and in other insects they are often found in the hemolymph and participate in antimicrobial defense. In *D. melanogaster*, however, rather than immune defense, lysozymes are strongly expressed in the digestive tract and are believed to be involved in the digestion of bacteria in the food (Daffre *et al*, 1994). Interestingly, ingestion of bacteria results in the induction of Nos in the digestive tract (Foley & O’Farrell, 2003), raising the possibility that the subsequent production of NO induces the expression of LysX and other lysozyme genes. Thus, in *Nos* mutant animals, *LysX* is downregulated as we see here (Fig. 1C). It remains unknown, however, why *LysX* expression is upregulated in the *Dg, Dys*, and *Syn1* mutants. Interestingly, in the analysis mentioned above in which we filter for genes dysregulated in *Dg, Dys*, and *Syn1* mutant genotypes (excluding the *Nos* LOF), two more lysozyme genes emerge, *LysE* and *LysS*, both of which are strongly upregulated in DGC mutants (Supp. Table 3). This result indicates that the DGC actively inhibits the expression of these lysosome genes by a Nos-independent mechanism, and it is consistent with our hypothesis that Nos is required for their upregulation in the gut.

To attain a clearer picture of the types of proteins encoded by the DGC-dependent genes, we clustered the gene products by annotated function and physical interactions using the STRING database (https://string-db.org; (Szklarczyk *et al*, 2021)). Direct and indirect physical interactions are shown as lines between nodes (genes), which we have assembled manually according to their annotated biological processes (Fig. 1D). The products of the genes that are dysregulated in all four DGC mutants are quite broad in function and associated biological process, suggesting that DGC deficiency has wide-ranging effects on the organism. Despite this, several distinct clusters emerged from our analysis, including reproduction, transcription regulation, and metabolism. The finding that genes involved in reproduction are dysregulated by DGC mutations is consistent with the male infertility exhibited by dystrophin-utrophin double mutant mice (Chen *et al*., 2017; Hernandez-Gonzalez *et al*., 2005).

### Temperature and Starvation stress result in distinct transcriptional changes

To investigate the genome-wide transcriptional effects of different stresses on adult flies, we determined the transcriptome-wide changes in gene expression of at least 2-fold in wildtype flies subjected to either temperature stress (33°C for 5 days) or starvation stress (4 days with yeast paste-only diet; see Materials and Methods). These conditions result in very different transcription expression profiles (Fig. 2A &2B). Temperature stress results in many more downregulated than upregulated genes (280 of 357 dysregulated genes are downregulated; 78.4%; Supp. Table 4). This is similar to the results of a recent study that analyzed the human stress response to passive exposure to environmental heat at the transcriptomic level (Bouchama *et al*, 2017). This study revealed that the heat-reprogrammed transcriptome was predominantly inhibitory and that the differentially expressed genes encoded proteins that function in stress-associated pathways such as proteostasis, energy metabolism, cell growth and proliferation, and cell death and survival. The transcriptomic changes also included mitochondrial dysfunction, altered protein synthesis, and reduced expression of genes related to immune function (Bouchama *et al*., 2017).

**Figure 2:**
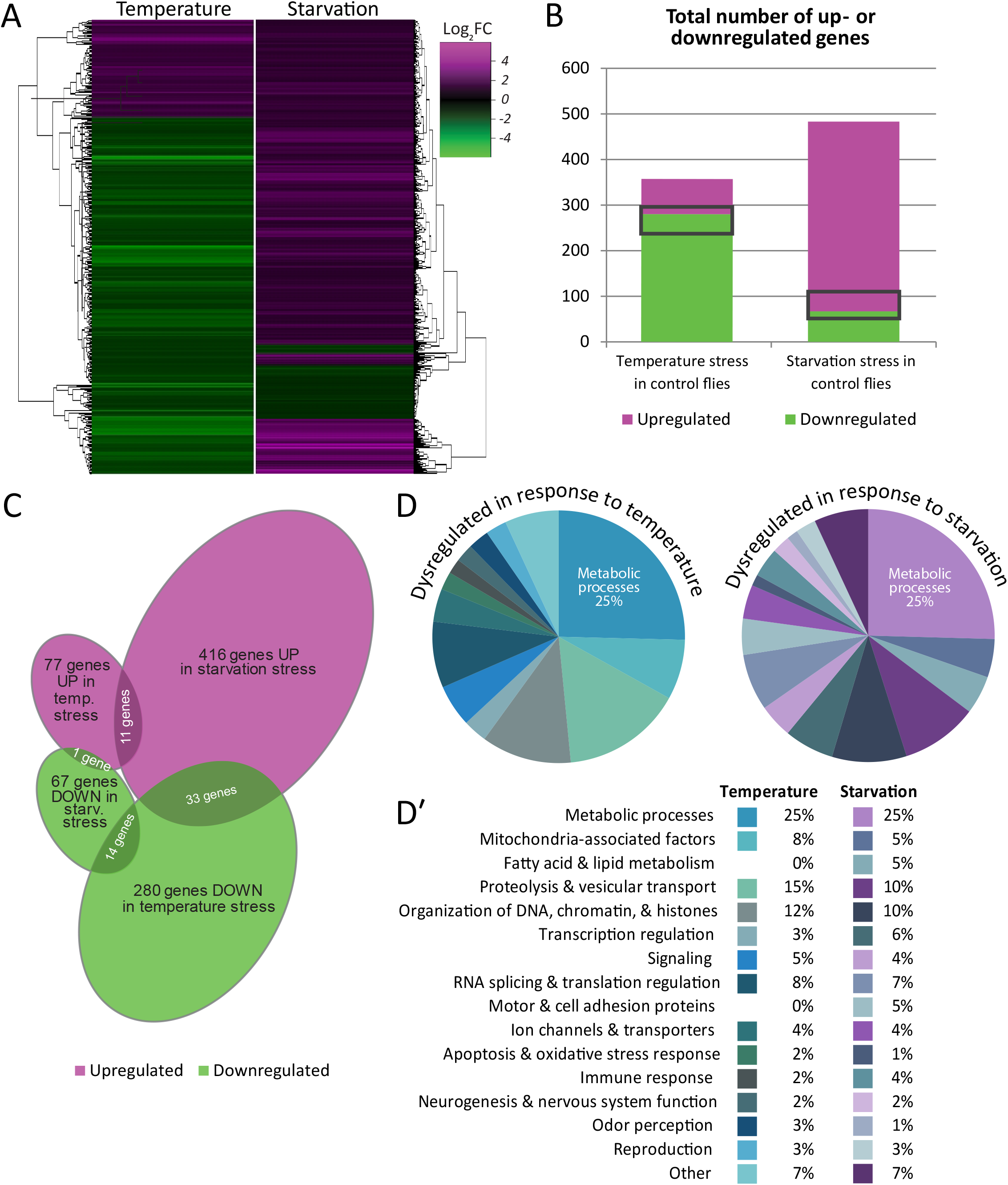
Temperature and Starvation stress result in distinct transcriptional changes (related to Supp. Fig. 2) A. Heat maps depicting all genes dysregulated in control flies by either temperature or starvation stress, irrespective of their levels of expression in the DGC mutants. B. Numbers of genes dysregulated in control flies by temperature stress (357 genes) or starvation stress (483 genes). Temperature stress is much more likely to cause downregulation of genes, while starvation stress causes more upregulation. Gray rectangles indicate the 59 genes that are dysregulated by both stressors. C. Venn diagram illustrating the up- and downregulation of genes as a result of temperature or starvation. The overlapping regions combined add up to the 59 genes dysregulated by both stressors. D-D′. Categories of all genes with known functions that are dysregulated in response to temperature or starvation. Pie charts (D) provide a visual depiction of the relative fraction each gene category comprises of the total. Table (D′) lists categories and percentages for both stress conditions. The largest category in both pie charts is Metabolic processes (25%), and the categories extend in a clockwise direction in the same order as in the table (D′).

In contrast to temperature stress, starvation stress is much more likely to cause upregulation (416 of 483 of dysregulated genes; 86.1%) (Fig. 2A & 2B; Supp. Table 5). The disparity we show between these different stress conditions is demonstrative of the fact that that temperature and starvation stress cause very different transcriptional outcomes. Heat shock protein (HSP) genes illustrate an important exception to this observation. Indeed, we found that of the 19 HSP genes we detected, most are weakly upregulated by temperature and downregulated by starvation stress (Supp. Table 1). Unexpectedly, however, none of these genes is upregulated more than 2-fold (log_2_FC>1) by temperature. We propose that this weak upregulation is due to the relatively mild and chronic heat treatment to which we subjected our flies. 16 out of 19 HSP genes are weakly upregulated by temperature with an average fold change of 1.3, while two genes are weakly downregulated. The final HSP gene, *Hsp23*, is the only HSP gene to show a dysregulation of more than 2-fold and exhibits a downregulation in temperature-stressed control flies (this downregulation is also exhibited by DGC mutants, placing *Hsp23* as a “DGC-independent” temperature stress-responsive gene; see below). Interestingly, a recent study demonstrated that even though *Hsp23* was upregulated by heat, its loss of function increased fruit flies’ tolerance to heat stress (Gu *et al*, 2021). It is possible that the intensity or duration of heat stress may affect the up- or downregulation of *Hsp23*; therefore, the temperature-responsive downregulation that we observe here may constitute part of a stress context-dependent survival mechanism.

Overall, a combined total of 840 genes are differentially expressed in temperature or starvation stress. Interestingly, the genes dysregulated in these two conditions exhibit little overlap: only 59 genes (7%) are differentially expressed in both conditions (Fig. 2B, gray rectangles; Supp. Table 6). Of these, 25 genes are similarly regulated – up- or downregulated in both stress conditions. The other 34 genes are differentially regulated, all but one being downregulated by temperature stress and upregulated in the starvation stress condition (Fig. 2C; Supp. Table 6). As temperature and starvation stress cause such disparate effects, it is possible that the 59 commonly dysregulated genes represent a general set of non-specific stress-responsive genes.

Next, we wanted to examine globally the types of genes that are dysregulated either by temperature stress or by starvation stress. We analyzed the list of dysregulated genes in each stress condition (Fig. 2A) and binned them according to their annotated functions. As 93% of these genes are dysregulated by only one stress condition but not the other, we expected to see that very different types and classes of genes would appear in each list. Surprisingly, we found that the contrary is true. Not only are similar classes of genes dysregulated by both stresses, the proportion assigned to each class is remarkably similar in both (Fig. 2D). For example, of all the dysregulated genes with annotated functions, metabolic genes comprise 25% of both temperature and starvation groups (Fig. 2D & 2D’). Similarly represented with very similar frequency in both temperature and starvation stress conditions were genes such as those encoding mitochondria-associated factors (8% in temperature stress; 5% in starvation stress), proteolysis-related genes (15% in temperature stress; 10% in starvation stress), transcription regulation (3% in temperature stress; 6% in starvation stress), as well as numerous other categories (Fig. 2D & 2D’).

Our finding that the most affected processes in response to high temperature were genes involved in metabolic regulation is consistent with previous studies. It has been shown that thermal stress affects metabolic and physiological functions and depletes energy reserves in *Drosophila* (Klepsatel *et al*, 2016; Klepsatel *et al*, 2019). Interestingly, a recent study demonstrated that the metabolism in stressed Drosophila depends on aerobic glycolysis – also known as the Warburg effect – rather than mitochondrial oxidative phosphorylation (Lee *et al*, 2015). Various genes involved in the electron transport chain and in ATP production were repressed, indicating poor mitochondrial fitness and suggesting that upon heat stress, there may be a switch in metabolism type in *Drosophila*.

Reproduction-related genes comprise 3% of genes dysregulated by temperature stress. It is not unexpected that reproduction is affected by thermal stress, since it has been reported that environmental stressors induce changes in endocrine state, leading to energy re-allocation from reproduction to survival (Meiselman *et al*, 2018; Ojima *et al*, 2018; Zwoinska *et al*, 2020). Stress-induced reproductive arrest has been documented not only in *Drosophila*, but also in other animals and humans (Boni, 2019).

Under starvation stress, the most highly represented category of dysregulated genes was ‘Metabolic processes.’ This group of genes was enriched for genes involved in aromatic amino acid metabolism and carbohydrate metabolism. The aromatic amino acids, phenylalanine, tryptophan, and tyrosine, are essential amino acids and are obtained in the diet, so the dysregulation of these genes and those involved in carbohydrate metabolism is consistent with a starvation state. We conclude that in response to stress, there is a common and reproducible set of processes that must be altered at the transcriptional level; while the specific genes dysregulated are stress-specific, the biological processes that are affected are common to different stresses.

### The transcriptional response to stress is defective in DGC mutants

Next, we sought to identify genes and pathways that do not respond appropriately to stress in the absence of a functional DGC. We began by examining our temperature-stress dataset for genes that exhibit a different response to temperature stress in control flies than in DGC mutants. First, we sought genes whose expression levels change in controls but do not change in any of the DGC mutants. For this purpose, we looked for genes which in controls have an expression change of at least 2-fold in response to temperature stress, and we considered genes to exhibit no change if their fold change was less than 1.62 (Log_2_<0.7). This analysis identifies 38 genes that fail to change expression levels in DGC mutants under temperature stress, even though they do change in controls (Fig. 3A; Supp. Table 7). We refer to this class of genes as “DGC-dependent response” genes, because their proper transcriptional changes in response to stress depend on an intact DGC. Complementarily, we also sorted for genes that do *not* exhibit an expression change in control (FC<1.62) but that change at least 2-fold in all DGC mutants. We found 8 genes with this pattern of expression in response to temperature; we refer to these as “DGC-prevented response” genes since their inappropriate expression changes (i.e., those that are not observed in control flies) are specifically prevented by the normal function of the DGC (Fig. 3A; Supp. Table 7).

**Figure 3:**
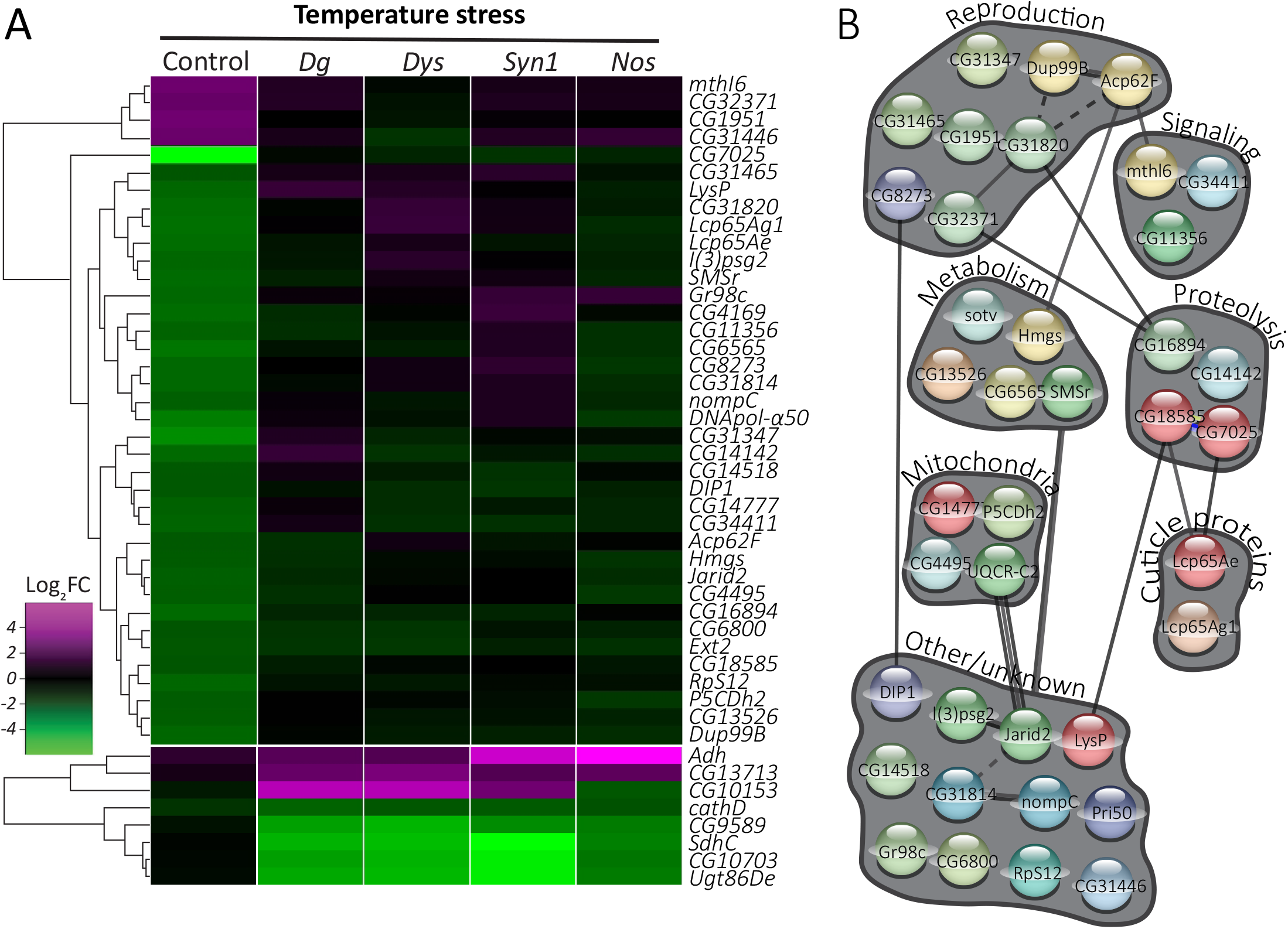
Defective transcriptional response to temperature stress in DGC mutants (related to Supp. Fig. 3) A. Heat map depicting DGC-dependent response genes (38 genes; temperature-responsive in control flies, but not in DGC mutants) and DGC-prevented response genes (8 genes; unchanged by temperature in controls, but with altered expression in DGC mutants). B. STRING-based clustering of the proteins encoded by the DGC-dependent genes depicted in A.

As it has been shown that MD patients have difficulty maintaining their body temperature (Hayes *et al*., 2008), this catalog of genes that do not respond appropriately to temperature may provide insight into the biological basis of this phenotype. In both DGC-dependent response and DGC-prevented response groups, dysregulated metabolic genes emerge (Supp. Table 7), indicating that DGC deficiency is detrimental to proper metabolism during stress, both by causing up- and downregulation of genes inappropriately and by failing to regulate genes that should be differentially expressed under stress.

Importantly, *mdx* mice are unable to maintain normal body temperature and have increased energy expenditure (Strakova *et al*, 2018). It has been speculated that *mdx* mice are not able to consume enough food to meet the metabolic demands of continuous muscle regeneration or that the thermoregulatory set point in the brain is defective in the absence of dystrophin. Similarly, Drosophila *Dg* mutants show abnormal metabolism (Kucherenko *et al*., 2011; Yatsenko *et al*, 2014a) and exhibit a cryophilic phenotype caused by increased energy (Takeuchi *et al*., 2009). This altered thermoregulatory behavior has been linked to the increased mitochondrial oxidative metabolism caused by activation of Ca^2+^ influx (Takeuchi *et al*., 2009). Loss of Dystrophin leads to abnormal, heat-sensitive muscle contractions that are repressed by mutations in Dys’s binding partner, Dystroglycan and can be rescued by blocking the Ca^2+^ channel (Marrone *et al*, 2011b). Moreover, *Dys* and *Dg* mutants have antagonistically abnormal cellular levels of reactive oxygen species (ROS), suggesting that the DGC has a function in regulation of muscle cell homeostasis (Kucherenko *et al*., 2011; Marrone *et al*., 2011b).

It has been shown previously that dystrophic muscles are already compromised, and as a consequence they are less adaptive and more sensitive to energetic stress and protein starvation (Kucherenko *et al*., 2011). Therefore, we were interested to identify genes that were differentially expressed in the DGC mutants upon dietary restriction. Applying the same analysis to genes dysregulated under starvation stress, we identified 61 DGC-dependent response genes and 4 DGC-prevented response genes (Fig. 4A; Supp. Table 8). Interestingly, none of the genes with perturbed DGC-dependent expression profiles are common to the temperature- and starvation-stress datasets. This underscores the distinct physiological reactions the organism manifests in response to the different stresses and suggests that a functional DGC is important in mounting both of these responses, albeit through different genetic pathways.

**Figure 4:**
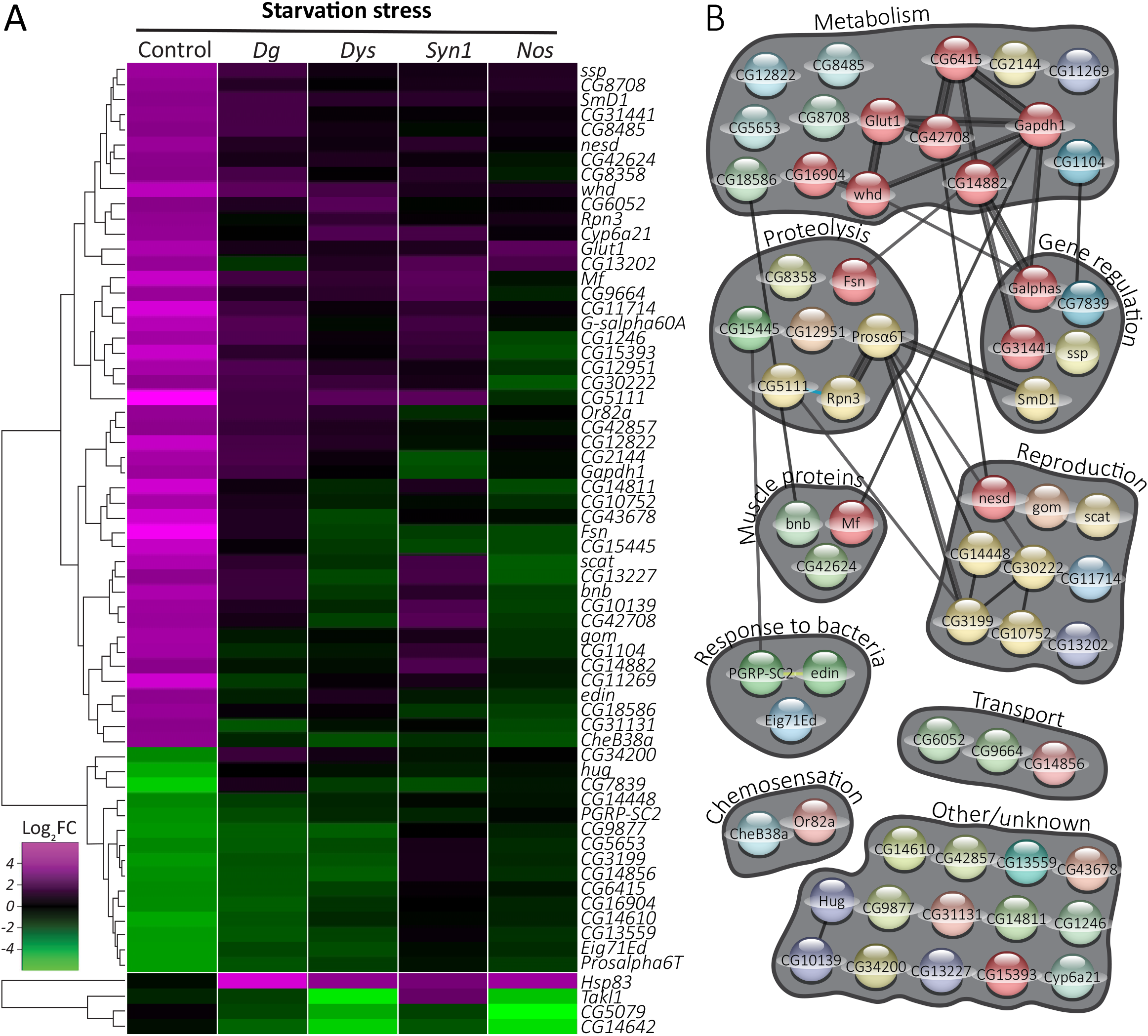
Defective transcriptional response to starvation stress in DGC mutants (related to Supp. Fig. 3) A. Heat map depicting DGC-dependent response genes (61 genes; starvation-responsive in control flies, but not in DGC mutants) and DGC-prevented response genes (4 genes; unchanged by starvation in controls, but with altered expression in DGC mutants). B. STRING-based clustering of the proteins encoded by the DGC-dependent genes depicted in A.

Regarding the starvation “DGC-dependent response” cluster, the most prominent clusters were a group of enzymes involved in metabolism regulation, proteasome proteins, and immune response factors (Fig. 4B; Supp. Table 8). Among the first group were nutrition-related proteins such as a fatty acid elongase (CG16904); a dehydrogenase (glyceraldehyde-3-phosphate dehydrogenase, Gapdh1) that binds NAD and regulates the glycolytic process and has biological roles in myoblast fusion, oxidation-reduction process, glucose metabolic process, and somatic muscle development; a Glucose transporter (Glut1); a palmitoyltransferase (*whd*) involved in oxidative stress and starvation; and a GTPase (G protein α s subunit, Gαs), which is predicted to enable G protein-coupled receptor binding activity and has been shown to positively regulate feeding behavior, response to trehalose, and sensory perception of sweet taste (Ueno *et al*, 2006). Moreover, similarly to *Dys* and *Dg* mutants, *Gαs*-deficient animals exhibit synaptic dysfunctions and have abnormal developmental rates (Hou *et al*, 2003; Renden & Broadie, 2003).

Interestingly, *whd* mutants exhibit an abnormal immune response, and there were other proteins involved in immune response in the DGC-dependent group. Edin is involved in the humoral immune response to Gram-negative bacteria. PGRP-SC2 (Peptidoglycan-recognition protein SC2) is an N-acetylmuramyl-L-alanine amidase that degrades biologically active bacterial peptidoglycans into biologically inactive fragments (Costechareyre *et al*, 2016; Guo *et al*, 2014). Recent studies of PGRP-SC2 mutants linked the immune and insulin signaling pathways. It has been shown that PGRP-SC2 downregulation produced InR-like phenotypes (Musselman *et al*, 2018).

Only four genes showed DGC pathway-prevented stress response: *Heat shock protein 83* (*Hsp83*, which encodes the only member of the HSP90 family of chaperone proteins in Drosophila) was significantly upregulated, while *Tak1-like 1* (*Takl1/TAK1*), *CG5079*, and *CG14642* were downregulated in starved mutants but not in starved controls (Fig. 4A).

Heat shock protein 83 (HSP83/HSP90) has been previously associated with response to various stresses in Drosophila and in mammals (Biebl & Buchner, 2019; Pearl, 2016). Interestingly, even in the absence of heat stress, the *Hsp83* gene is expressed at high levels in multiple tissues during development (Xiao & Lis, 1989; Zimmerman *et al*, 1983). HSP90 is a molecular chaperone that promotes the maturation, structural maintenance, and proper regulation of specific target proteins involved for instance in cell cycle control and signal transduction (Bandura *et al*, 2013). HSP90 employs the energy of ATP hydrolysis to control the folding and activation of client proteins (Mader *et al*, 2020). In addition, it dynamically interacts with various co-chaperones that modulate its substrate recognition, ATPase cycle and chaperone function. For example, in muscles, Hsp90 binds myosin via a scaffold protein Unc-45 (Ni & Odunuga, 2015). This interaction is required for proper myosin folding and protection from stress (Bujalowski *et al*, 2018; Lee *et al*, 2011). Interestingly, in the presence of dietary amino acids, Hsp90 is both necessary and sufficient for neuronal stem cell reactivation by promoting the activation of InR pathway in the developing brain (Huang & Wang, 2018).

In the DGC-dependent groups for both starvation and temperature stress, there are prominent clusters of metabolic, proteolysis-related, and reproduction-related (primarily testis-expressed) genes. Most of the genes in these groups follow the general trend in which genes are downregulated upon temperature stress and upregulated under starvation. For example, 100% of the DGC-dependent metabolic genes (*Ext2, CG6565, Hmgs, SMSr*, and *CG13526*) and the proteolysis-related genes (*CG16894, CG18585, CG7025*, and *CG14142*) in the temperature stress group are downregulated in controls, while in the starvation condition, most (75%) of the metabolic genes (*CG1104, CG14882, CG18586, CG42708, Gapdh1*, and *Glut1*) and 80% of the proteolysis genes (*CG15445, CG5111, Fsn*, and *Rpn3*) are upregulated.

The DGC-dependent temperature- and starvation-stress-dependent genes that are involved in reproduction (Fig. 3B & 4B), along with those reproduction-related genes that are DGC-dependent even in unstressed conditions (Fig. 1D), could present important new insight into the molecular causes of fertility deficits in DGC-deficient mice and flies.

In addition to these similarities, the DGC-dependent groups also differ in important ways between temperature and starvation. In temperature stress, there is a group of mitochondrial genes that are downregulated; no such group appears in starvation. Similarly, in starvation stress, there is a group of regulatory genes (G proteins, transcription factors, splicing factors), mostly upregulated in controls, while no such group appears in temperature stress.

### A subset of genes exhibits DGC-independent responses to temperature and starvation

Interestingly, while generating the DGC-dependent and DGC-prevented response gene datasets, we found almost no genes that are regulated in response to stress in an entirely opposite way in control vs. DGC mutants (i.e., upregulated in control but down in all the mutants, or vice versa). Indeed, if we filter the temperature and starvation stress datasets such that we can examine all genes that are similarly regulated in the four DGC mutants, we find that there are three categories of genes: 1) changed in controls but not in mutants (DGC-dependent stress response genes); 2) unchanged in control but altered expression in all mutants (DGC-prevented stress response genes); and 3) changed in controls *and* in DGC mutants. Nearly all of the genes in the latter category are regulated the same direction, up or down, in all genotypes, including control (Supp. Fig. 3). Overall, these genes respond to temperature stress irrespective of genotype; therefore, we consider them to represent “DGC-independent response” genes.

In the cluster of genes that were changed in all genotypes in response to temperature stress, 90% of genes (19 of 21) were regulated in the same direction (up- or down-regulated) (Supp. Fig. 3A; Supp. Table 9). That feature was not a criterion for inclusion in our analysis; rather, we required was that a gene was 2-fold dysregulated in all genotypes, but any gene could have been up in some genotypes and down in others, and still been included here. But almost all genes are dysregulated in the same direction in all genotypes. The two exceptions are the outliers *bigmax* and *Brd*. Each is downregulated by temperature in all except one genotype: *bigmax* is upregulated in the *Dys* mutant, and *Brd* is upregulated in the *Dg* mutant.

Similarly, in the starvation-stress cluster, 88% of genes (37 of 42) followed this pattern, corresponding to a DGC-independent response (Supp. Fig. 3B; Supp. Table 10). Of the few outliers, most exhibited differential regulation between the DGC mutants. The sole exception was *sro*, which, in response to starvation stress, is upregulated in control and downregulated in all of the DGC mutants (Supp. Fig. 3B; Supp. Table 10).

Several lines of evidence in flies and mammals have established the DGC as a signaling hub. First, the association of Nos with Dys establishes the DGC as an important component of the conserved nitric oxide signaling pathway. Syn-associated Nos produces NO, which is used in the nitrosylation of intracellular proteins. The nitrosylation of histone deacetylases (HDACs) affects the expression of downstream genes, including those encoding microRNAs, which can have a negative-feedback role on the expression of Dg (Cacchiarelli *et al*., 2010; Yatsenko *et al*., 2014b). In addition, the DGC has also been shown to physically associate with Yorkie (Yki) and Kibra (Kbr), components of the Hippo signaling pathway, affecting the expression of downstream genes (Morikawa *et al*., 2017; Yatsenko *et al*., 2020). Finally, murine Dg and insulin receptor (IR) are closely associated in the muscle sarcolemma. In aging muscle, Dg undergoes increased internalization and degradation via the lysosome, and due to the proteins’ association, IR levels are also reduced. The reduction of IR in aged muscle can be expected to have far-reaching transcriptional effects due to disruption of the insulin signaling pathway. Indeed, this mechanism may help to explain the insulin insensitivity phenotype exhibited by dystrophic patients (Eid Mutlak *et al*., 2020). The finding in this study of a cluster of transcriptional regulators whose expression is regulated by the DGC (Fig. 1) implies that the transcriptional effects exerted by the DGC may be much more far-reaching than was previously known. Further research will determine whether the transcriptional changes in DGC mutants occur only through these pathways or if other signaling and transcriptional pathways are involved.

## Materials and Methods

### Fly Stocks

All fly stocks and crosses were maintained at 25°C on a standard cornmeal-agar-based food in a 12-12h light-dark cycle. Fly stocks used in this study (see also Table 1) were *w*^*1118*^, *Canton-S, w*^*1118*^*;Df(2L)BSC230/CyO*, and *w*^*1118*^*;Df(3L)BSC450/TM6C,Sb*^*1*^*cu*^*1*^ from the Bloomington Drosophila Stock Center (BDSC); *Dg*^*O55*^*/CyO* and *Dg*^*O86*^*/CyO* (kind gifts from Robert Ray); outcrossed *Df(3R)Exel6184* deficiency (Marrone *et al*., 2012); *Nos*^*Δ15*^ (kind gift from P. O’Farrell); and *ΔSyn1*^*5-2*^*/TM3* (see below).

*ΔSyn1* mutants were generated using two transgenic lines, *w*^*1118*^; *PBac[WH]CG14565*^*f05859*^ (BDSC #18911) and *P[XP]CG7370*^*d06092*^ (Exelixis at Harvard Medical School) (Thibault *et al*, 2004), containing transposon elements with FRT sites flanking the *Syn1* gene. In addition, *w*^*1118*^ *hsFlp; Dr/TM3,Sb* was used for heat shock-induced Flp recombinase expression. The recombination event was induced by heat shocking embryos and early larvae (0-36h collection) with the genotype *w*^*1118*^ *hsFlp; PBac[WH]CG14565*^*f05859*^*/P[XP]CG7370*^*d06092*^ in a 37°C water bath three times for 1h each in 12h intervals. Emerging flies with mosaic eye color were crossed to *w*^*1118*^; *Ly/TM3,Sb* and their progeny were selected for the white-eye trait as an indicator of loss of mini-white gene cassettes from both transgenes, which indicates completion of the recombination event, i.e. the deletion in the *Syn1* locus (Supp. Fig. 1A) (Parks *et al*, 2004). Multiple *ΔSyn1*^*5-2*^ alleles were isolated independently using this procedure and molecularly validated as described below. The isolate *ΔSyn1*^*5-2*^ was used in trans over a deficiency as the *Syn1* LOF in all subsequent experiments in this study.

*ΔSyn1* alleles have a ∼26kb long deletion in the 3^rd^ chromosome removing the *Syn1* gene, confirmed by two PCRs (Supp. Fig. 1B and 1B′) by HotStarTaq Master Mix (Qiagen) using the following primers: CAATCAACATGAAGAGCCAACCCA (*Syn1*-Exon-5-Fw) and ACTTTGCCGCCGATGTCACTGT (*Syn1*-Exon-5-Rv) to confirm the absence of the *Syn1* gene (Supp. Fig. 1B, red arrows), AATGATTCGCAGTGGAAGGCT (XP5′ plus), GACGCATGATTATCTTTTACGTGAC (WH5′ minus) (Parks *et al*., 2004) to detect the recombination-resulted junction of the two residual transgene fragments (Supp. Fig. 1B′, red arrows). In addition, an absence of *Syn1* mRNA expression in *ΔSyn1* mutants was confirmed by qPCR using High Capacity Reverse Transcriptase (Applied Biosystems) and Fast SYBR Green reagents in a StepOne Plus Real Time PCR System (Applied Biosystems) using the following primers for *Syn1* and *RpL32*, respectively: CCCTCGTCTGGTTCAATGCC (*Syn1*-Fw), AATCTCAAATACATCGACCC (*Syn1*-Rv), AAGATGACCATCCGCCCAGC (*RpL32*-Fw), GTCGATACCCTTGGGCTTGC (*RpL32*-Rv). For the calculation of relative mRNA expression, the housekeeping gene *RpL32* C_T_ values were subtracted from the C_T_ values of *Syn1* to calculate the ΔC_T_ values. The *Syn1* expression in *Oregon-R-C* flies was analyzed as control and for normalization, hence calculation of the ΔΔC_T_ values. Relative expression was calculated with the formula 2^-ΔΔCT^ (Supp. Fig. 1C).

### RNA sequencing and data analysis

Samples were prepared for RNA sequencing as follows. One-week-old flies were used with the following genotypes: *Canton S/w*^*1118*^ (*Control*), *Dg*^*086*^*/Dg*^*055*^ (*Dg LOF*), *Df(3R)Exel6184/ Df(3R)Exel6184* (*Dys LOF*), *Df(3L)BSC450/ΔSyn1*^*5-2*^ (*Syn1 LOF*), and *Df(2L)BSC230/Nos*^*Δ15*^ (*Nos LOF*). Flies were kept on standard food for 5 days at 25°C in a 12h light/dark cycle for the unstressed condition. In order to induce heat stress, flies were kept in an incubator at 33°C on standard food for 5 days. Starvation stress was induced by feeding the flies on yeast paste only (yeast paste made from dry yeast mixed with 5% propionic acid in water) at 25°C for 4 days.

For RNA extraction, ∼10 male flies per genotype and condition were homogenized together in Trizol (Ambion), and RNA was extracted using Direct-zol RNA mini-prep with an additional on-column DNAse digestion step (Zymo Research). The quality of the purified RNA was assessed with a Nanodrop ND-1000 spectrophotometer measuring the A260/A280 and A260/A230 ratios. Only RNA with A260/A280>2.0 and A260/A230>1.7 was used for further RNA-seq application. For each sample, 5-10 µg of total RNA was sent to GATC Biotech AG (Konstanz, Germany) for library preparation and subsequent transcriptome sequencing. In summary, RNA-seq libraries were prepared by RNA poly-A purification, fragmentation, random primed cDNA synthesis, linker ligation, and PCR enrichment. The samples were used to make a random-primed cDNA library, and the run was performed on an Illumina HiSeq platform with single-end, 100bp reads.

The transcriptome sequencing experiment resulted in a sample average of ∼7.5 million reads that could be mapped to a unique transcript. Genome index was generated from genome FASTA files of individual chromosomes (BDGP6 version) and the transcript annotation GTF file Drosophila_melanogaster.BDGP6.84 (ensemble.org database (Yates *et al*, 2016)). Subsequently, the reads were mapped to the reference genome using STAR: ultrafast universal RNA-seq aligner (Dobin *et al*, 2013). Subsequent analyses were performed in R statistical software (Team, 2016) using packages from the Bioconductor project (Gentleman *et al*, 2004). Resulting BAM alignment files were used to generate counts on individual transcripts using transcript database from the same GTF file (via Rsamtools (Morgan *et al*, 2010) and GenomicFeatures (Lawrence *et al*, 2013) packages) and “summarizeOverlaps” function in “Union” mode via GenomicAlignments package (Lawrence *et al*., 2013). In total, 15930 unique transcripts were detected with at least 1 count. Next, the counts were analyzed and genotype and condition comparisons were done through built-in statistical models in the DESeq2 package (Love *et al*, 2014). For data visualization, gplots (Warnes *et al*, 2013) and RColorBrewer (Neuwirth, 2011) packages were used.

For differential gene expression analysis, 2-fold difference and p values smaller than 0.1 were considered significant and filtered by the “results” function with “lfcthreshold = 1” and subsetting the genes with p value <0.1. The resulting gene lists were subjected to gene interaction and ontology term analysis using STRING and DAVID databases and ClueGO (Bindea *et al*, 2009) and CluePedia applications via Cytoscape software (Shannon *et al*, 2003).

## Acknowledgements

We would like to thank Nai-Hua Hsiao and April Marrone for the generation of the *Syn1* deletion mutant; all members of the Shcherbata lab for insights and comments on the manuscript; Marko Shcherbatyy for drawing of DGC scheme; VolkswagenStiftung, MPIBPC, and MHH for funding; GATC Biotech AG for sequencing; BDSC and Exelixis at Harvard Medical School for fly stocks.

## Supplementary Figure Legends

**Figure 1:**
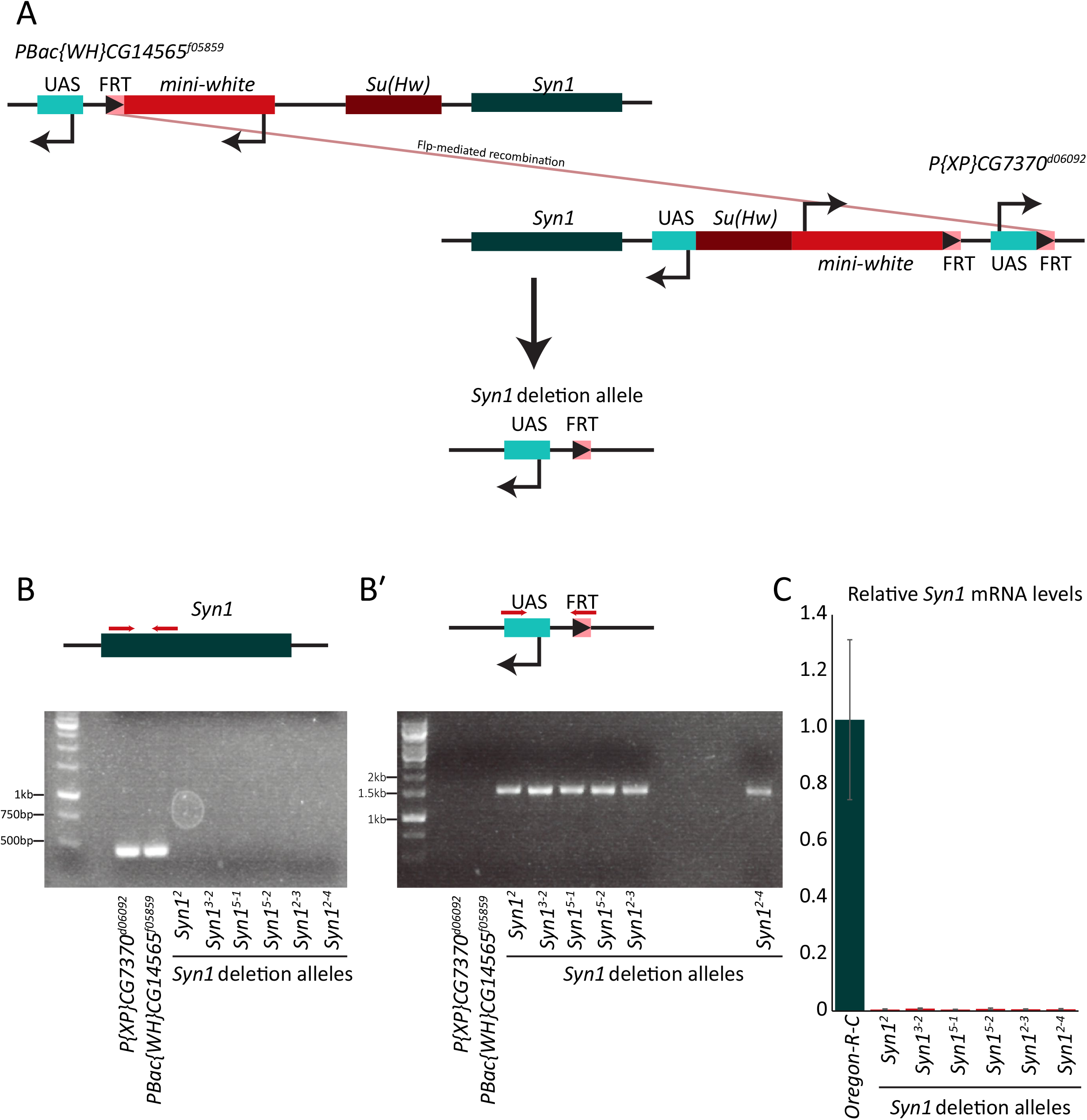
Generation of new *Syntrophin1* deficiency, *ΔSyn1*. A. Deficiency was generated using Flp-mediated recombination between two FRT-containing transposons flanking the *Syn1* gene (*PBac[WH]CG14565*^*f05859*^ and *P[XP]CG7370*^*d06092*^; see Materials and Methods). B-B′. PCR confirmation of new *ΔSyn1* alleles. (B) Primers specific for the endogenous *Syn1* gene result in PCR amplification in the parental strains *PBac[WH]CG14565*^*f05859*^ and *P[XP]CG7370*^*d06092*^, but no bands are present in lanes representing *Syn1* deletions. (B′) Primers that recognize residual transposon fragments flanking the new deficiency positively detect the presence of the deficiency in flies with deletion alleles, but not in parental flies. C. RT-qPCR confirms the absence of RNA from *Syn1* deletion mutants, while it is readily detected in control flies (Oregon-R-C).

**Supplementary Figure 2:**
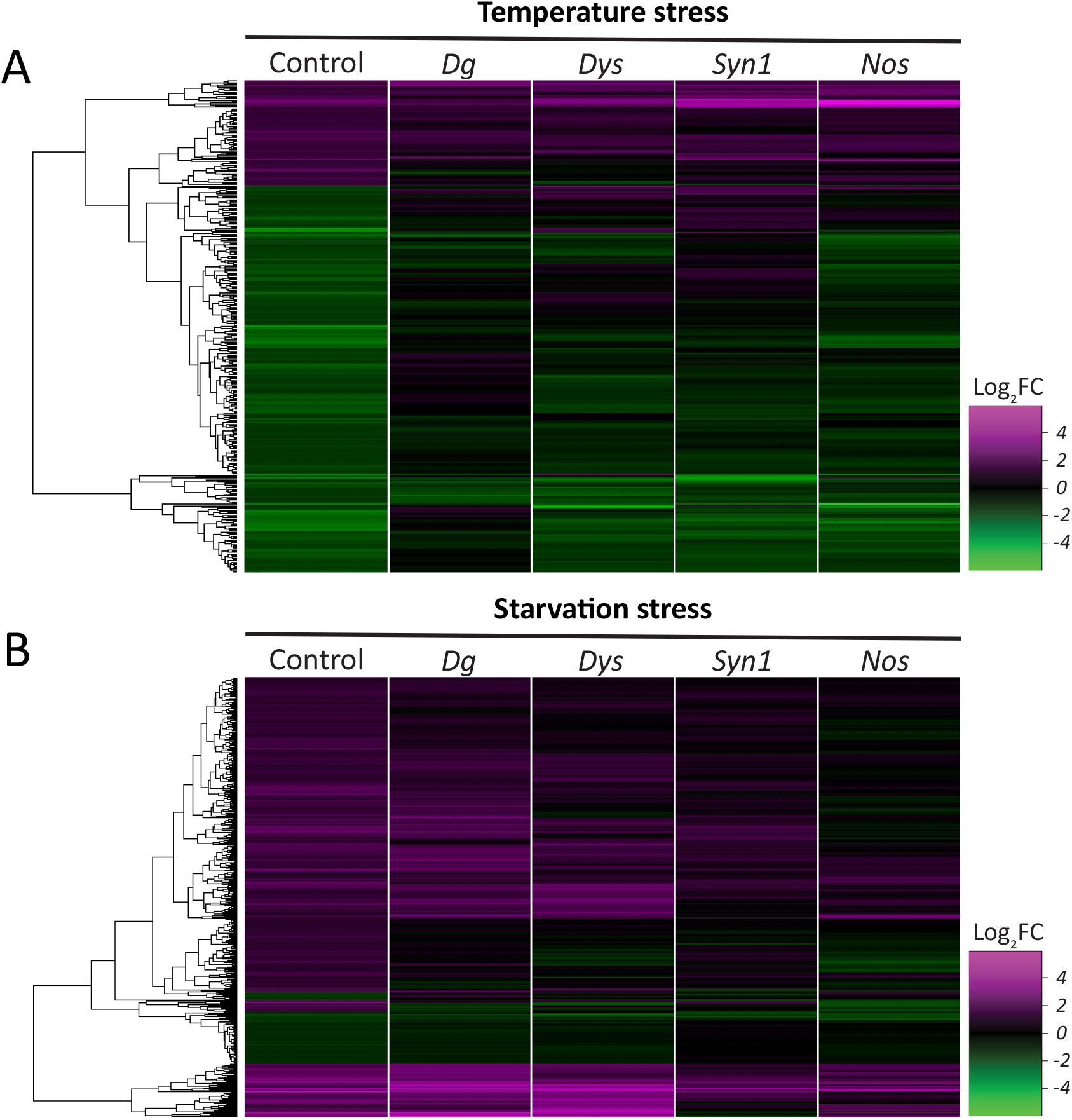
All genes dysregulated by temperature and starvation stress (related to Figure 2) A-B. Heat maps with all genes dysregulated in control flies by temperature stress (A; 357 genes) and starvation stress (B; 483 genes), also showing the expression level of these genes in the four DGC mutants.

**Supplementary Figure 3:**
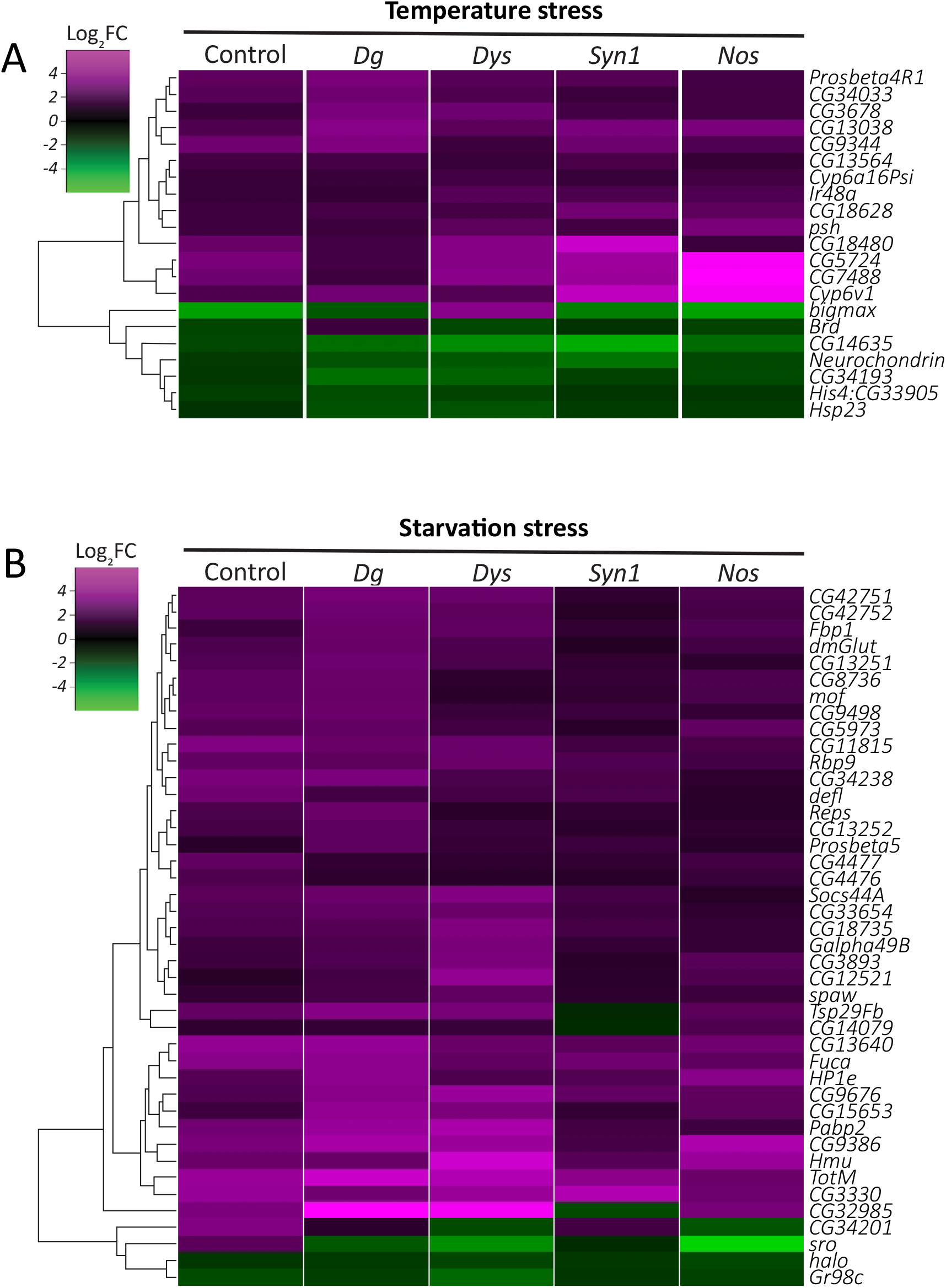
DGC-independent genes are regulated by stress irrespective of presence of a functional DGC (related to Figures 3 and 4) A-B. Heat maps showing all genes that are dysregulated by temperature stress (A; 21 genes) or starvation stress (B; 42 genes) in all genotypes, including controls, indicating a DGC-independent stress-response mechanism. Of these genes, 90% of temperature-responsive genes and 88% of starvation-responsive genes are dysregulated similarly (up- or downregulated) in all genotypes.

